# Reproducing Left Ventricular Twist by Mimicking Myocardial Fiber Orientation Using 3D Bioprinting-assisted Tissue Assembly

**DOI:** 10.1101/2023.07.03.547595

**Authors:** Dong Gyu Hwang, Hwanyong Choi, Uijung Yong, Donghwan Kim, Wonok Kang, Sung-Min Park, Jinah Jang

## Abstract

**Background:** Left ventricular twist, an opposite rotation of the apex and base, is caused by myocardial fiber orientation, a unique structural feature of the myocardium, and contributes to the effective ejection fraction of the native heart. Reproducing this structural-functional relationship in an *in vitro* heart model remains challenging due to the lack of synchrony between layers when mimicking the fiber orientations of each layer.

**Methods:** We employed a hierarchical approach for creating multilayered and multiaxial fibers in a chamber-like structure, as follows: 3D bioprinting-assisted tissue assembly, fabrication of uniaxially aligned engineered heart tissue as a building block, and assembly of them to create a myocardial fiber orientation in a chamber-like structure.

**Result:** The EHT module confirmed uniaxial alignment and cardiac functions such as contractility and electrophysiological properties. By fabricating the assembly platform by 3D bioprinting, it is possible to guide building blocks in various directions as intended, confirming the versatility of this method. The assembly platform allows structural and functional synchrony of assembled tissues while controlling and maintaining predefined cellular alignment. Furthermore, various shapes and sizes of EHT modules and assembly platform were fabricated for mimicking myocardial fiber orientation in a chamber-like structure. The resulting structure exhibited three layers and three orientations representing myocardial fiber orientation. Moreover, the left ventricular twist was confirmed by measuring basal and apical rotations.

**Conclusions:** Recapitulation of the microscale structure of the left ventricle enabled the identification of information not discernible from the existing macroscale structure. This understanding of the structure-function relationship of the heart can provide insights into the mechanisms underlying cardiac structure, function, and related diseases. Furthermore, the versatility of the 3D bioprinting-assisted tissue assembly allows for the creation of organs and tissue collections with complex structural and functional features by fabricating and assembling modules that meet the specific requirements of target tissues and organs.

## Introduction

The left ventricle (LV) has the thickest myocardium of any heart chamber and produces contractile forces that allow for blood circulation throughout the body. Microscopically, uniaxially aligned cardiomyocytes (CMs) generate contractile forces, and the myocardial fibers composed of CMs have unique orientations that gradually change from the epicardium to the endocardium (Fig. 1A, i). Myocardial fibers that are helically aligned in opposite directions in the subepicardial and subendocardial regions contract in each direction, contributing to LV twist (LVT), in which the apex and base rotate in opposite directions, thereby promoting effective blood circulation ^1–3^. Thus, numerous efforts have been made to induce and control cellular alignment in engineered heart tissues (EHTs) to understand and implement the structure-function relationship of the LV. Traditional approaches for inducing cellular alignment include seeding cells onto predefined geometrical cues ^4–10^, encapsulating cells within extracellular matrix (ECM)-based hydrogels, and molding them into grasping structures ^11–17^. Although these methods are effective, it is difficult to create multiaxial alignment throughout different layers. Therefore, a hierarchical approach — stacking ^18^ or wrapping ^19^ a highly aligned structure to achieve three different orientations ranging from -60° to +60° in different layers — was introduced to recapitulate myocardial fiber orientation (MFO). Nonetheless, prior approaches continue to lack structural scalability, and the presence of additional scaffolds hinders between-layer coupling.

**Fig. 1.**
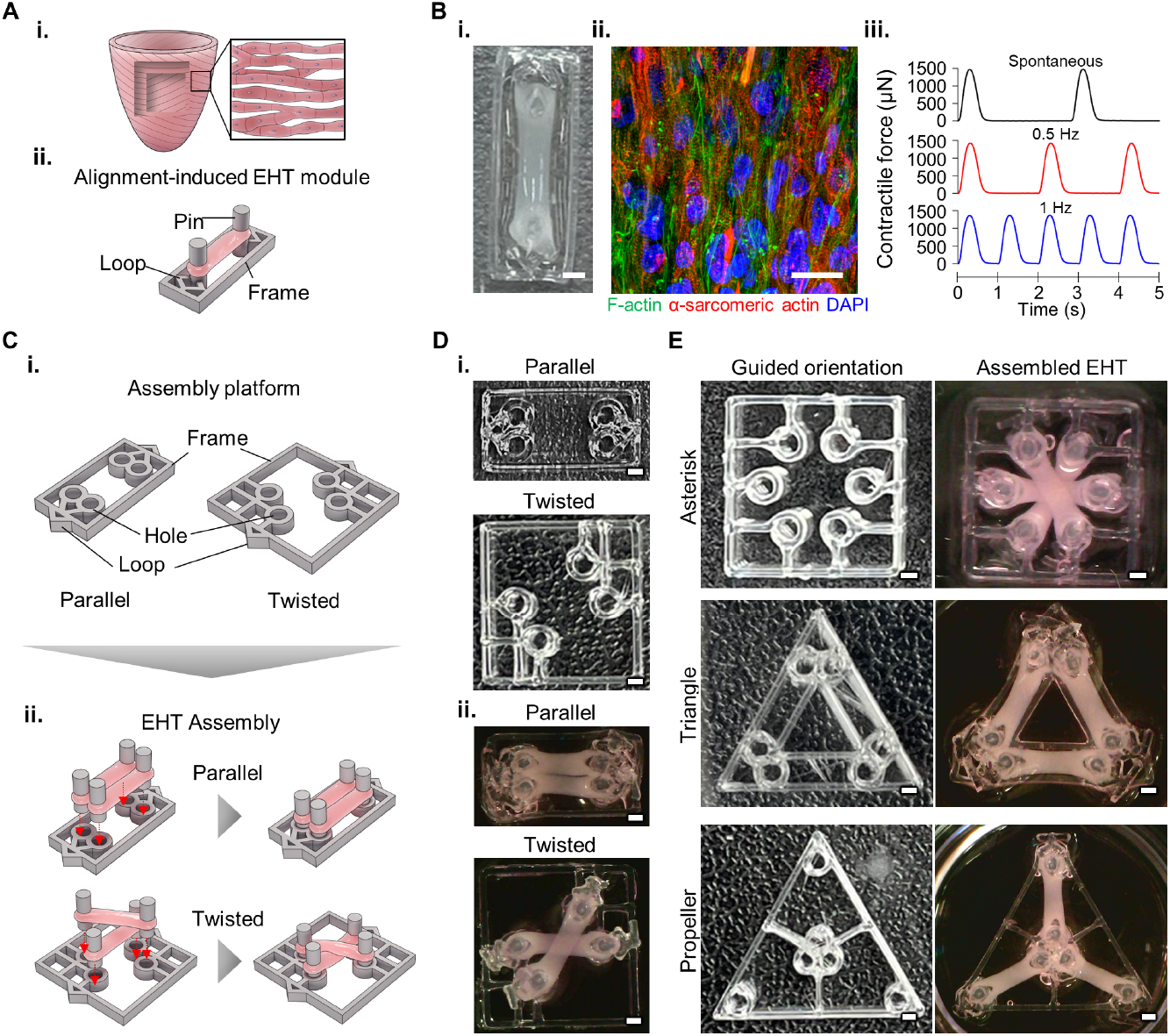
Versatility of the 3D bioprinting-assisted tissue assembly strategy. Schematic representations of (**A**) a myocardial fiber (i) and an engineered heart tissue (EHT) module (ii) as a functional and structural unit; and a (**B**) 3D-bioprinted EHT module (i; scale bar, 1 mm) exhibiting uniaxially aligned cellular orientation (ii; green, F-actin; red, α-sarcomeric actin; blue, 4′,6-diamidino-2-phenylindole [DAPI]; scale bar, 25 μm) and contractility (iii; top, spontaneous; middle, 0.5 Hz paced, bottom, 1 Hz paced). (**C**) A schematic diagram of two types of EHT assembly platforms (EAPs) (i) and the assembly process (ii); (**D**) an image of 3D-printed EAPs, (i) with two assembly types using two EHT modules (scale bars, 1 mm); and (**E**) the rapid prototyping of an EAP for tri-assembly, with assembled EHTs created using tree modules (scale bars, 1 mm).

Three-dimensional (3D) biofabrication is gaining popularity owing to its high structural reproducibility, which facilitates the generation of LV models and even multichambered shapes that closely resemble the architecture of the native heart ^20^. However, cell alignment has not yet been incorporated into bioprinted chamber models. Recently, a circumferentially and helically aligned ventricle model was fabricated using jet-spinning fiber technology, demonstrating the relationship between structure and cardiac function and presenting apical rotation in a helical model ^21^. Although this approach generated a dual-chambered fiber structure mimicking native ECM alignment, cells were distributed only on the surface and not in a transmural manner after seeding. Given that the native LV contracts simultaneously throughout the tissue, it is necessary to induce between-layer coupling of structure and function of myocardial fibers in addition to mimicking complex alignment directions to reproduce LVT.

We previously developed a cardiac tissue-specific bioink to enhance cell function; however, we encountered low printing fidelity due to its low viscosity ^14,17,22–24^. To address this issue, we incorporated light-activating molecules into the bioink to facilitate the creation of tissue constructs of high complexity with fidelity ^25,26^. Nevertheless, our attempts to modulate the decellularized extracellular matrix (dECM) bioink did not allow us to avoid the inherent trade-off between biological and fabrication properties. A distinct strategy was required for reaching resolution smaller than the diameter of the printing nozzle ^27,28^. Hence, we created 3D bioprinting-assisted tissue assembly (3DBT), a hierarchical approach for fabricating both micro- and macroscopic structures of the LV while harnessing the superior biological properties of tissue-specific bioink to reproduce LVT.

The approach involved the utilization of 3D bioprinting technology for tissue assembly, enabling the versatile manufacturing of necessary elements such as modules and assembly platforms. This strategy consisted of the following two main steps: manufacturing uniaxially aligned EHT modules and tissue assembly to control their multiaxial alignment. As building blocks, EHT modules reproduce the structural (i.e., uniaxial cellular alignment) and functional (i.e., contraction and electrophysiological properties) features of myocardial fibers. Tissue assembly processes and platforms were used to control and build EHT modules to drive functional and structural coupling in a single tissue. Furthermore, it was confirmed that the versatile manufacturing of the EHT module and assembly platform allowed the structure and function of the module to be controlled in the desired direction and shape. We used this strategy to reproduce LVT by creating MFO in a chamber-like structure (MoCha). MoCha, composed of three layers with right-handed helical, circumferential, and left-handed helical orientations from the deep to the outer layer. Moreover, the between-layer coupling that has not been achieved before was induced by interlocking the EHT modules, demonstrating LV motions such as multiaxial deformation and LVT. In summary, we reproduced the complex structural and functional features of the LV by assembling uniaxially aligned EHT modules into multiaxially aligned MFO, leading to the transformation of uniaxial cardiac tissue contraction into LV motions, including LV deformation and twisting.

## Methods

The authors declare that all supporting data and methodological descriptions are available within the supplemental materials. Please also see the Major Resources Table in the supplemental materials.

## Results

To achieve uniaxial cellular alignment, the EHT module consisted of polymer and tissue parts (Fig. 1A). We fabricated the EHT module by printing a polyethylene vinyl acetate (PEVA) structure containing two posts capable of imparting tensile stress, followed by printing the cell-laden bioink onto the structure (fig. S1 and movie S1). CMs and stromal cells (e.g., fibroblasts and endothelial cells) have been extensively used to match the physiological relevance of EHTs to that of native tissues ^29^. Cardiac fibroblasts (CFs) are recognized for their involvement in ECM remodeling and their facilitation of functional connectivity between CMs ^30^. Indeed, we confirmed that the mixture of human induced pluripotent stem cell-derived CMs (iPSC-CMs) and CFs in the bioink (CM + CF) enhanced cellular alignment compared to the encapsulation of CMs only (CM) (Fig. 1B, i-ii, S2, and movie S2). In addition, uniaxial alignment was sufficiently achieved 5 days after EHT generation (fig. S3). EHT modules exhibited synchronized and spontaneous contractility along with pacing in response to electrical stimulation, indicating that they had essential cardiac functionality (Fig. 1B, iii and movie S3).

Recent advances in tissue assembly have enabled the construction of 3D tissue structures with enhanced structural complexity by locating spheroid building blocks in desired locations ^31–34^. However, these approaches are inadequate for creating a directional construct such as an MFO. In fact, an assembly strategy using anisotropic building blocks in a controlled manner has not yet been reported ^35,36^. To guide the uniaxially aligned EHTs to their desired positions and angles, we developed an EHT assembly platform (EAP). Inspiration for this design was taken from LEGO^®^ bricks in which a few simple building block shapes can be assembled via interlocking parts to create a variety of complex structures. We employed a concave-convex structure that allows for LEGO^®^ bricks to be connected; therefore, the EAP was designed and fabricated to include a concave (i.e., hole) structure capable of connecting to the convex (i.e., post) structure of the EHT module (fig. S4 and movie S4). On this basis, the versatile manufacturing of EAPs, especially due to the locations of holes, facilitated controlling the orientation and location of the EHT modules (Fig. 1, C-D). Furthermore, the versatility of this strategy was demonstrated by creating asterisk, triangle, and propeller structures using three EHT modules (Fig. 1E).

Functional synchronization, structural coupling, and orientation control of the EHT module are required for the construction of an MFO using tissue assembly. To investigate whether these requirements were satisfied using the EAP, we employed parallel (pa-EHT) and twisted (ta-EHT) assemblies. We first confirmed that two individually functioning EHT modules showed synchronized beating, with the border between the modules blurred after 6 hours of assembly (movie S5). Furthermore, the two distinct EHT modules gradually fused as new tissues formed between them (Fig. 2A and S5), with the degree of fusion quantified by calculating the fusion index (Fig. 2B and S6). To clarify why synchrony and fusion were observed, assemblies were stained to identify α-sarcomeric actin (α-SA), vimentin (Vim), and connexin 43 (Cx43). Immunofluorescence images and cross-sectional fluorescence intensity profiles of the assemblies showed that Cx43, a gap junction marker, was highly expressed at tissue interfaces, indicating junction formation between the two modules. In addition, the expression pattern of α-SA, a marker for CMs, was similar to that of Cx43, indicating that functional synchronization was achieved by the junctional formation of iPSC-CMs. Vim expression surrounded the outside of the EHT modules and was predominantly present in newly formed areas, suggesting that the CFs contributed to the formation of new tissues at the fusion interface (Fig. 2C, and S7C). These findings indicate that self-assembly of cells occurs during functional synchronization and structural coupling. However, during this self-assembly process, cells may not only be rearranged within each module but also migrate to adjacent modules, resulting in unintended changes in the existing cellular alignment ^32,37^. Therefore, we conducted in-depth analyses of cellular self-assembly at the interface of EHT modules to identify any changes in alignment during the fusion process. To investigate whether cell migration occurs and to determine what cell types are involved in self-assembly, two types of EHTs were assembled, unlabeled, and labeled with Dil and DiO solutions for iPSC-CMs and CFs, respectively. The ratio of the labeled-region fluorescence to the unlabeled-region fluorescence of the two tissues revealed no significant change in the fluorescence signal or width ratio of the two tissues after 10 days (Fig. 2D, and movie S6). Further image analysis of the EHTs after disassembly indicated that both cell types were exclusively at the module interface. In addition, a cross-sectional view of the unlabeled EHT using confocal imaging showed that cells had migrated to the newly formed area rather than inside the unlabeled EHT (fig. S7, A-B). These results suggest that the cells maintained their original territories rather than actively travelling to other tissues. We sought to confirm the influence of cell alignment during the self-assembly process by measuring the mean resultant vector length (MRVL) distribution. Consequently, pa-EHT showed uniaxially aligned cells in the main body as well as the fusion area, whereas ta-EHT presented bidirectional alignment following programmed orientation (Fig. 2E, i-ii and S8). Moreover, there were no significant changes in the module or MRVL after assembly, and no between-group differences in MRVL were observed (Fig. 2E, iii). These findings indicate that our strategy can be used to regulate cellular alignment such that cells are placed in their intended orientations without significantly affecting the predefined alignment while inducing functional and structural coupling.

**Fig. 2.**
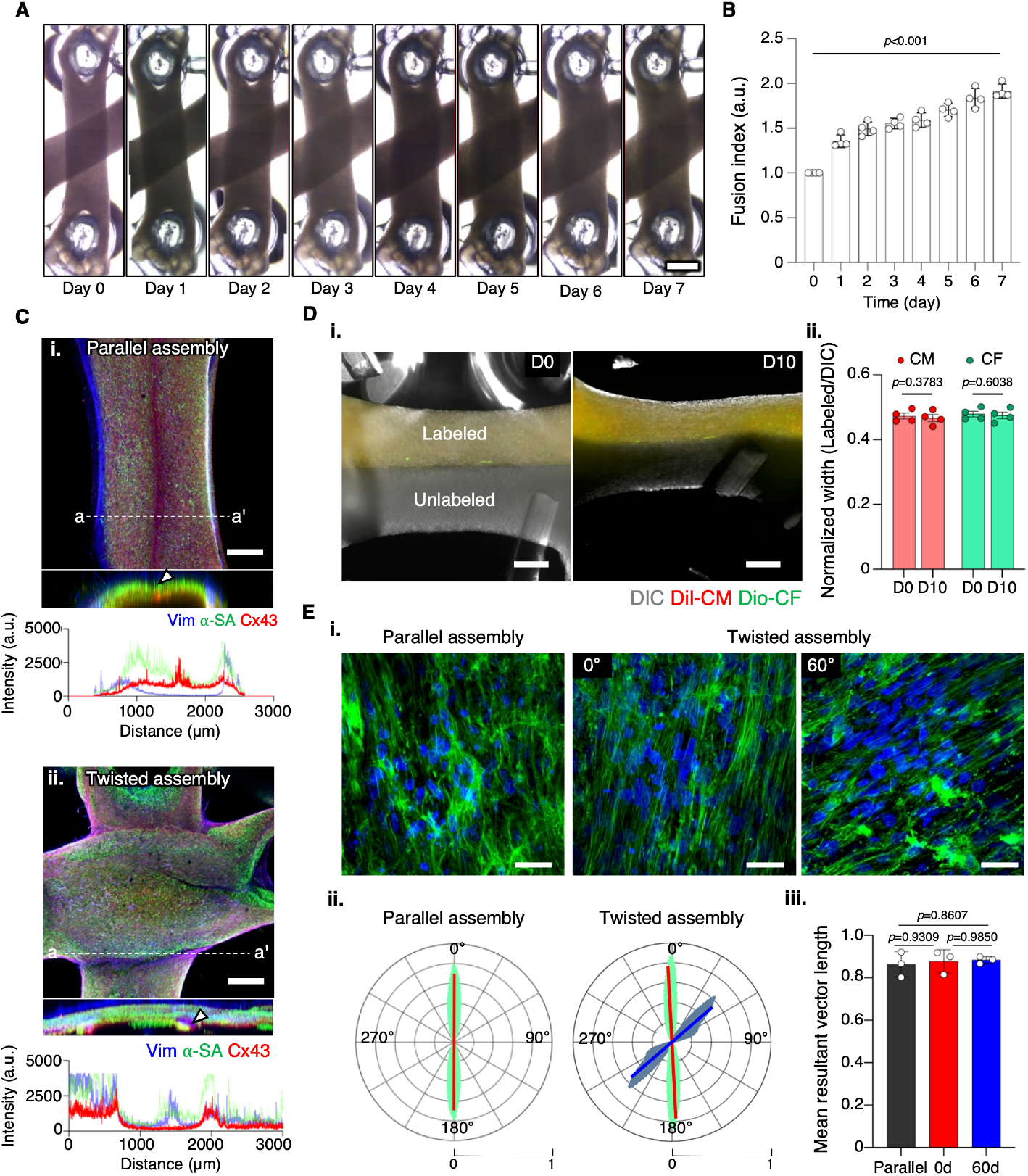
Induction of functional and structural coupling using engineered heart tissue (EHT) assembly platforms. (**A**) Stitched images of the twisted assembly, in chronological order, showing newly formed tissues (scale bar, 1 mm). (**B**) Quantification of changes based on newly formed area (*n* = 4, *p-*value via multiple comparisons with day 0, using one-way analysis of variance [ANOVA]). (**C**) Fluorescent images of parallel (i) and twisted (ii) assemblies showing structural coupling and fluorescence signal intensity profiles and expression levels of cardiac cell markers and junction proteins (a-a’, cross-sectional area; blue, vimentin; green, α-sarcomeric actin; red, connexin 43; scale bars, 500 μm). (**D**) Assessment of cell migration using fluorescent images of parallel assembly with labeled and unlabeled modules at days 0 and 10 (i; gray, differential interference contrast (DIC); red, Dil-labeled cardiomyocytes; green, Dio-labeled cardiac fibroblasts; scale bars, 500 μm) and quantitative data by normalizing each fluorescence signal width with the DIC width (ii, n = 4 by Šídák’s two-way ANOVA). (**E**) Assessment of cellular orientation according to assembly. Representative fluorescent images of each group (i; scale bars, 25 μm), and quantitative data indicating cellular orientation distribution (ii) and mean resultant vector (iii).

Given the pivotal role of the structural-functional relationship in the native LV, we sought to determine whether controlling structural orientation using EAPs allows us to control the functional orientation of the structure. The contraction motion was analyzed using particle imaging velocimetry (PIV), with orientation and velocity distributions calculated (movie S7). The contraction motion and velocity of pa-EHT were mainly observed in the longitudinal and transverse cellular alignment directions, which corresponded to the EHT module, whereas those of ta-EHT were observed in the circumferential direction (Fig. 3, A-B, and S9A). In addition, results of the MRVL calculation showed that the larger vector was in the pa-EHT in which the alignment was concentrated in one direction, whereas the smaller vector was in the ta-EHT owing to its dispersed contraction motion. (Fig. 3C and S9B). Although our initial expectation was that the contraction motion of a single module and parallel assembly would follow a longitudinal direction consistent with their cellular alignment, the actual contraction motions were captured transverse to the alignment. This discrepancy may be attributed to the isometric contraction that occurred because both ends were fixed to the PEVA contraction. Therefore, we conclude that PIV analysis identified the vector in accordance with wall thickening ^20^. Contractile motion changes associated with assembly direction were also confirmed based on contractile force generation. The contractile force generated along the longitudinal direction of the EHTs was measured (fig. S10A). The contractile forces of pa-EHT and ta-EHT were approximately 1.5 times and 1.2 times those of the EHT module, respectively, indicating that actual values tended to mirror those that were predicted in which the module produced a force F, and pa-EHT and ta-EHT generated forces of 2F and 1.5F, respectively (Fig. 3, D-E, and S10B). However, we confirmed that there were no significant changes in contraction time (e.g., time to peak and 90% relaxation time), implying that structural modulation using EAP only affected the contractile force (fig. S10C). Subsequently, we investigated the action potential (AP) propagation characteristics associated with the assembly direction using microscopic analysis. As a result, optical mapping of each group showed that the AP propagated throughout each tissue, convincing us that assembled EHTs functioned as a single tissue. Moreover, we observed transmural propagation of the AP, starting from a junctional point in the ta-EHT, followed by alignment of the part at a 60° angle from the ta-EHT (Fig. 3F and movie S8). Conduction velocity was analyzed in the longitudinal (LCV) and transverse (TCV) directions, directions of cell alignment ^21^. The LCV and TCV of pa-EHT were comparable to those of the EHT module, indicating that propagation properties were maintained while cellular alignment was preserved. Similarly, AP propagation of ta-EHT followed cellular alignment (Fig. 3, G-H).

**Fig. 3.**
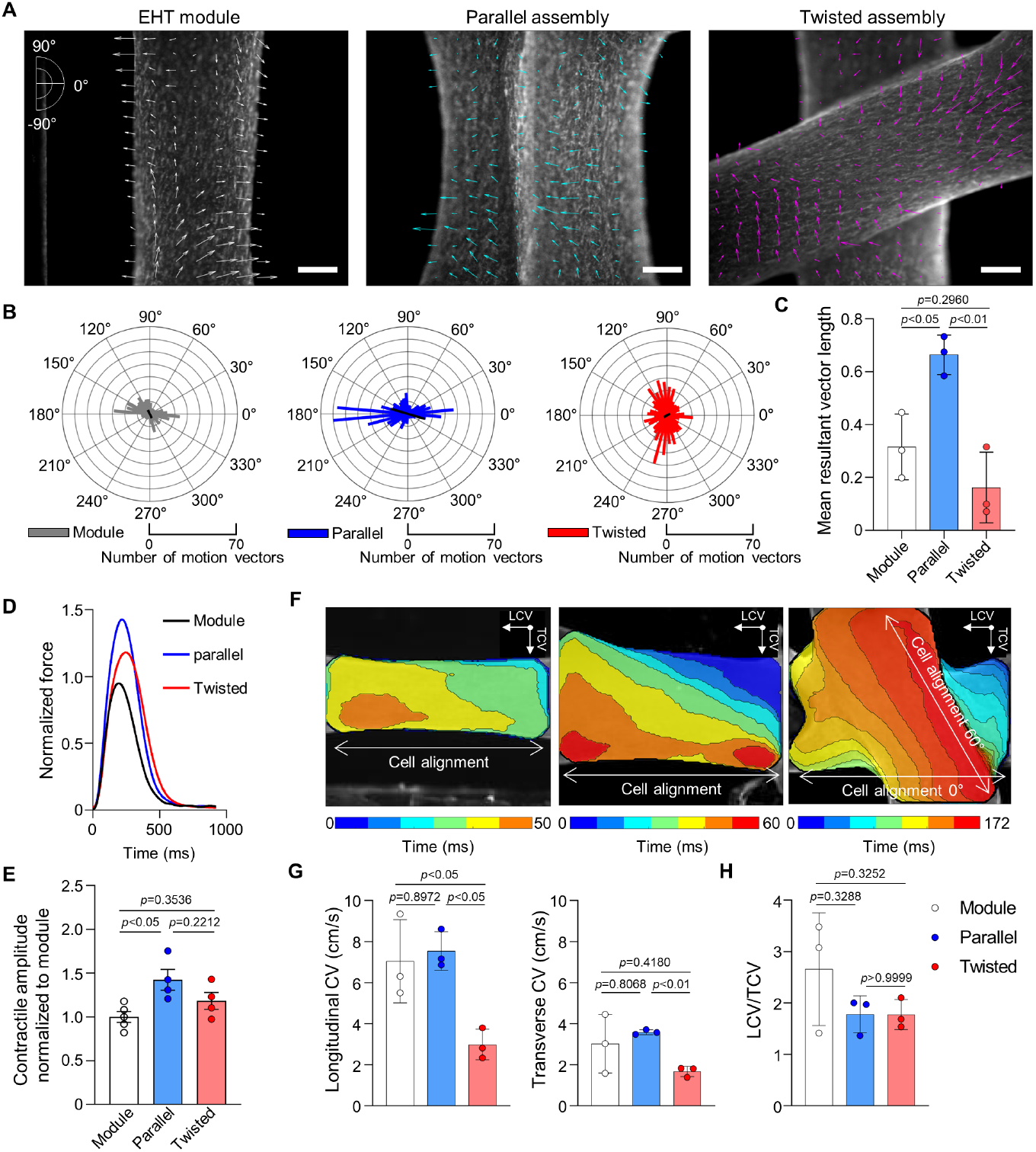
Functional changes according to assembly orientation. (**A**-**C**) Assessment of contraction direction after assembly. (A) Representative images used for particle imaging velocimetry (PIV) analysis (scale bars, 500 μm); (B) a polar histogram showing the cumulative distribution of motion vectors for all samples used in the PIV analysis, and (**C**) Quantitative data of mean resultant vector length of motion vectors. (**D** and **E**) Contractile changes after assembly are shown using (D) mean contractility and (E) quantitative data indicating normalized contractile amplitudes (n = 5, module; n = 4, parallel and twisted). (**F**) Optical mapping of action potential propagation shows synchronized signal propagation through the tissues (scale bars, 500 μm). (**G**) Calculation of conduction velocity along the longitudinal (LCV) and transverse (TCV) directions of cellular alignment allows for the determination of propagation using LCV/TCV ratio.

We confirmed that 3DBT can induce the functional synchrony and structural coupling of assembled aligned modules and that the orientation of cardiac functions can be controlled after assembly. On this basis, we envisioned that 3DBT is able to facilitate the manufacturing and assembly of EHT modules to build the MoCha, leading to the reproduction of LV-like motion (Fig. 4A) ^2,3,38^. To manufacture the MoCha, we fabricated longitudinally aligned, strip-type EHT modules (s-EHTs) for deep and outer layer configurations, and circumferentially aligned, ring-type EHT modules (r-EHTs) to construct middle-layer configurations (Fig. 4B and S11). Moreover, an EAP was created to guide EHT modules into a chamber-like structure. The EAP consisted of two parts and was designed to guide module placement in the desired orientation and geometry and to induce tissue fusion. The geometry-guiding part was designed to be removed for observation after inducing sufficient tissue fusion (Fig. 4C). Tilted holes of the orientation-guiding part allowed s-EHTs to form right- and left-handed helixes. To create the MoCha, one end of the s-EHT was anchored to a deep layer to form a right-handed helix. In the middle layer, r-EHTs were placed on top of the right helixes to form fibers in a horizontal orientation. Finally, opposite ends of s-EHTs were fastened to form a left-handed helix to create an outer layer (Fig. 4D and S12). The MoCha contained gradual orientation changes of EHT modules from the deep layer to the outer layer, with interlocking spirals at its apex, mimicking the structural features of the LV (Fig. 4E and S13A) ^39^. Synchronization of the global structure was confirmed by measuring the beat of each MoCha layer (Fig. 4F). Thereafter, we sought to demonstrate that the structure and function of MoCha can reproduce those of the LV. Chamber-like structures exhibit different deformation behaviors depending on the orientation of their fibers. Circumferentially aligned fibers contribute to deformation primarily in the basal region, whereas helically aligned fibers result in deformation of the apical region ^21^. We assessed MoCha deformation by measuring percentage area change in the long-axis view. Consequently, MoCha, which included all MFOs, exhibited deformation in both the basal and apical regions, with the apical region displaying greater deformation than the basal region (fig. S13B). Furthermore, we evaluated the rotation of the MoCha by tracking one point in the basal and apical regions in the short-axis view. Strikingly, results revealed clockwise rotation of the base and counterclockwise rotation of the apex. Further, the rotation angle of the apex was greater than that of the base, a finding in agreement with observations of the long axis. LVT angle was determined by calculating the difference in the rotation angle between the two regions, revealing that rotation angle patterns observed were consistent with those of actual clinical measurement (Fig. 4G and movie S9) ^2^. Although the LVT observed in this model is feeble compared to the native LVT ^21^, it can offer new opportunity to implement cardiac structure-function relationships, just as the earlier chamber-like heart models provided insight into the cardiac volume-pressure relationship ^2,40,41^.

**Fig. 4.**
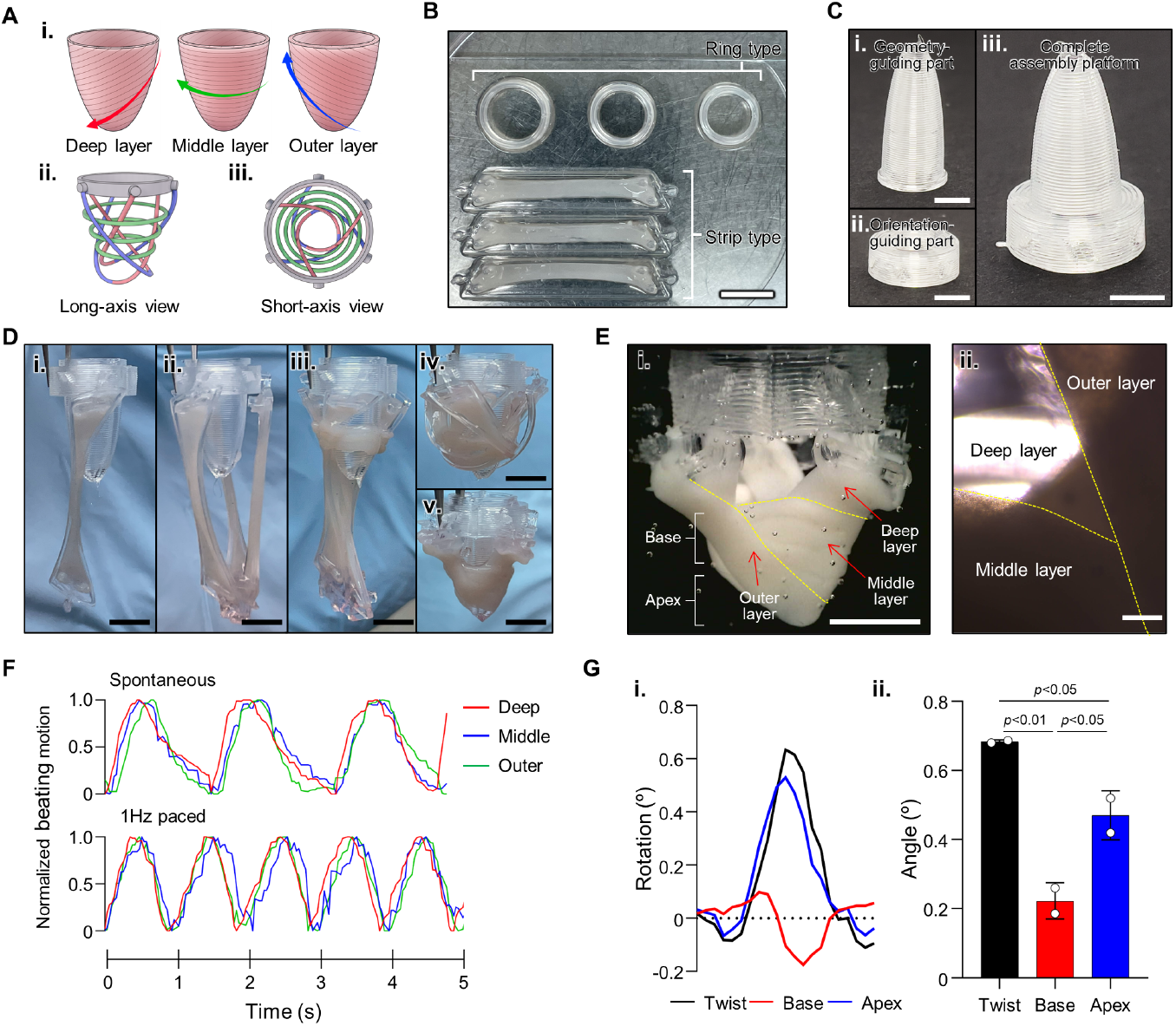
Reproduction of LV motion via mimicking myocardial fiber orientation in a chamber-like structure. (**A**) Schematic illustration of (i) three layers showing the orientation of the native left ventricle and (ii) the assembly strategy used to create the model (red, deep layer with right-handed helix; green, middle layer with circumferential fibers; blue, outer layer with left-handed helix). (**B**) Various shapes and sizes of EHT modules are used for myocardial fiber orientation (MFO) in a chamber-like structure (MoCha) assembly (scale bar, 1 cm). An **(C)** assembly platform consisting of orientation-(i) and geometry-(ii) guiding parts, and a (iii) final construct after assembly of the parts (scale bars, 5 mm). **(D)** Photographs depict the MoCha assembly process. (I and ii) Fixation of one end of s-EHT forms a right-handed helix in the deep layer, (iii) r-EHTs are placed over the deep layer to form circumferentially aligned fibers, (iv) fixation of another end of s-EHTs forms left-handed helix in the outer layer, and (v) frame removal is performed at the final stage of assembly (scale bars, 5 mm). **(E)** A (i) photograph of the whole MoCha structure after disassembly of a bullet-like structure (scale bar, 5 mm) and (ii) microscopic images presenting the transmural orientation of three layers (scale bar, 500 μm. **(F)** Measuring the beating motion of the three MoCha layers indicates that they beat in a synchronized manner. **(G)** Left ventricle twist in a short-axis view from its base, with (i) representative graphs showing basal and apical rotation and a left ventricle twist (positive, counterclockwise; negative, clockwise) and (ii) the quantification of rotation angles (n = 2).

## Discussion

3DBT provides a hierarchical approach for realizing the complex structural and functional features of the LV that have not yet been achieved, despite considerable effort. By assembling one-dimensional modules to replicate a native three-dimensional MFO, the one-dimensional contraction of these modules can be transformed into three-dimensional deformation and rotation. In particular, the first LVT-mimicking construct provided insights into the mechanisms underlying cardiac structure, function, and related diseases. For instance, this construct can serve as a drug safety and efficacy testing platform *in vitro* by exhibiting changes in LVT. Furthermore, the versatility of the 3DBT approach allows for the creation of organs and tissue collections with complex structural and functional features by fabricating and assembling modules that meet the specific requirements of target tissues and organs.

## Article Information

### Author contributions

D.G.H. and J.J. designed the study. D.G.H. and J.J. wrote and revised the manuscript. D.G.H., U.Y., and D.K. designed and fabricated EHT modules and EAP. D.G.H. performed functional validation of tissue modules and assembled tissues. D.G.H. and H.C. conducted and analyzed the EP study. D.G.H., U.Y., D.K., and H.C. cultured cells and tissues. W.K. and S.-M.P. manufactured the electrical stimulation system. J.J. supervised the research. All authors discussed the results and contributed to the writing of the final manuscript.

### Materials and Data Availability

All data are available in the main text or the supplementary materials.

## Non-standard Abbreviations and Acronyms

3D: Three dimensional
3DBT: Three dimensional bioprinting-assisted tissue assembly
AP: Action potential
α-SA: α-sarcomeric actin
CF: Cardiac fibroblast
CM: Cardiomyocyte
Cx43: Connexin 43
dECM: Decellularized extracellular matrix
EAP: EHT assembly platform
ECM: Extracellular matrix
EHT: Engineered heart tissue
iPSC-CM: Induced pluripotent stem cell-derived cardiomyocyte
LCV: Longitudinal conduction velocity
LV: Left ventricle
LVT: Left ventricular twist
MFO: Myocardial fiber orientation
MRVL: Mean resultant vector length
MoCha: Myocardial fiber orientation in a chamber-like structure
PEVA: Polyethylene vinyl acetate
PIV: Particle imaging velocimetry
pa-EHT: Parallel-assembly EHT
r-EHT: Ring-type EHT
s-EHT: Strip-type EHT
TCV: Transverse conduction velocity
ta-EHT: Twisted-assembly EHT
Vim: Vimentin

## Supplementary Materials

Materials and Methods

Figs. S1 to S13

Table S1

References (*42-44*)

Movies S1 to S9

Major Resources Table

